# A New Large-Scale Extracellular Vesicle Production Strategy for Biomedical Drug Development

**DOI:** 10.1101/2024.04.15.589541

**Authors:** Shiyi Yang, Lei Zhang, Xin Zhou, Xincheng Peng, Xin Zhang, Wenli Wang, Jinxiu Zhao, Xinjun He, Ke Xu

## Abstract

Extracellular Vesicles (EVs) are nanoscale to microscale membranous vesicles released by cells. They are vital for intercellular communications in physiological and pathological conditions, offering immense potential in diagnostics and therapeutical applications due to their ability to transfer diverse biological cargos between cells. While research of EVs is continuously gaining popularity in academia, large-scale manufacturing aiming for clinical applications remains challenging. Herein, we introduce a novel large-scale EV production strategy, including an upstream process with fed-batch suspension HEK293 cell culture, and a downstream process with chromatographic purification for ultra-pure EVs. Such promising EV production strategy enables the potential capability of large scale GMP manufacture of EVs for clinical applications.

## 2. Introduction

Extracellular vesicles (EVs) have garnered significant attention in recent years owing to their diverse roles in intercellular communication and their potential applications in various fields, particularly in biomedicine. These nanosized membranous vesicles, released by almost all cell types, play crucial roles in intercellular communication by facilitating the transfer of biomolecules including proteins, lipids, DNA, RNA, and miRNA between cells, enabling the exchange of genetic information, signaling molecules, and cellular components.^1,2^ Through these mechanisms, EVs participate in diverse physiological processes such as immune modulation, tissue regeneration, and neural communication. Moreover, EVs have been implicated in pathological conditions including Alzheimer’s disease,^3, 4^cancers,^5-7^and cardiovascular diseases,^8^highlighting their potential as diagnostic markers and therapeutic agents.^9^

The translation of EVs from basic research to clinical applications necessitates the development of scalable and reproducible manufacturing processes. While the exploration of EVs is burgeoning in academic research, scaling up production for industrial use remains a daunting task. Selecting the parent cell is one of the most crucial step. The characteristics and contents of EVs depend on the type of originating parental cells and their secretion status.^1^ A large portion of research focuses on human cell-derived EVs, such as those found in blood or plasma,^10, 11^ primary mesenchymal stem cells (MSCs),^12^ induced pluripotent stem cells,^13^ immortalized primary cells (such as MSC-MYC),^14^ or established cell lines like HEK293 (a human embryonic kidney cell line).^15^ There is also significant interest in EVs derived from non-human sources for therapeutic purposes, including microbial cultures,^16^ bovine milk-derived EVs,^17, 18^ plants,^19^ and marine microalgae.^20^ These alternative sources of EVs may offer broad availability and low cost. However, due to immunogenicity concerns, the therapeutic applications of these sources have to be carefully tested. Human cell derived EVs, due to their low immunogenicity in the human body, are preferred as drug delivery systems in biopharmaceutical development. Derived from human embryonic kidney cells transduced with adenovirus 5 (Ad5) DNA, the human HEK293 cell line possesses immortalization properties. HEK293 cell lines are easily grown in serum-free suspension cultures and are readily transfected for gene expression.^21^ One of their most important attributes is their human origin, making them more suitable for producing biologics for human use. Since its creation, this cell line has been extensively characterized and applied in the fields of recombinant proteins, cell therapy, Adeno-Associated Virus (AAV)-based biopharmaceutical drug development, and other areas. Compared to stem cells, establishing large-scale culture in chemical defined medium for HEK293 cell lines is relatively easier. These properties make HEK293 the good candidate for generation of a large quantity cells and uniform EVs without altering their phenotype.^22^ Additionally, genetic modification of HEK293 cells is relatively straightforward, facilitating engineered modifications to express EVs with specific information.^23^

The purification of extracellular vesicles (EVs) from complex biological samples is a critical step in their isolation and characterization. However, conventional lab-scale EV purification methods are not ideal for large-scale downstream processes due to various limitations. For example, Ultracentrifugation is the most widely used EV isolation method in academic labs, known as the gold standard of the EV purification approach.^24-27^ It separated the EVs based on differential centrifugal forces. However, the whole process is time consuming, and the yield is undesirable.^24, 27^ Also, a lack of purity of isolated EV is a problem due to co-pallet of impurities during conventional UC. Density gradient ultracentrifugation enhances EV purity by layering the sample onto a density gradient medium by adding sucrose or iodixanol.^28, 29^ However, its thin loading layer make it unsuitable for large-scale applications.^30^ Polymer-based precipitation method utilizes polymers (e.g. polyethylene glycol) to induce the aggregation and precipitation of EVs from solutions, such as cell culture medium or clinical blood.^31^ But this approach suffers from insufficient purity due to co-precipitation of impurities and the difficulty of the polymer removal.^32, 33^ Ultrafiltration is a prominent frequently used method for the purification of EVs.^34^ Several small and large scale membranes with different molecular weight cut-off are commercially available for the removal of impurities from EV with different sizes. However, EV loss occurs due to membrane adhesion and clog, and mechanic driving force may cause deformation and damage to EVs.^25^ Immuno-affinity capture utilizes antibodies targeting specific EV surface markers to selectively isolate EVs. While offering high purity, this method’s scalability is limited by antibody availability and the complexity of large-scale processing. At the current stage, there are still significant challenges in large-scale purification of EVs to support preclinical and clinical research or commercial markets.

Several pioneering biotech companies have achieved significant milestones in the mass production of extracellular vesicles (EVs). One such company, Codiak Biosciences, emerged as a clinical-stage biopharmaceutical leader in EV-based drug development. By leveraging a 2000-liter bioreactor, Codiak successfully scaled up their upstream production processes. Moreover, their downstream processing team utilized a combination of filtration techniques such as ultrafiltration/diafiltration (UF/DF) and liquid chromatographic processes including cation exchange chromatography (CEX), anion exchange chromatography (AEX) for EV purification.^35^ Another notable player in the field is EVOX Therapeutics, a biotechnology company specializing in EV-based therapies for rare diseases such as Phenylketonuria, cardiovascular diseases, and neurological disorders.^36^ They have achieved their upstream capability with 500 L perfusion bioreactor, and they claim the ability to further scale up their production to 2000 L. Their EVs are purified with filtration processes and liquid chromatography.^37^ Leveraging scalable manufacturing processes to produce clinical-grade EVs, EVOX engineered their EV products to deliver diverse drug payload, including AAV vectors, genome editing modalities, and RNA therapeutics.

As a pre-clinical stage biotechnology company focusing on developing innovative EV-based therapeutics for regenerative medicine and cancer therapy, herein we introduce a new massive EV production strategy in this study. The upstream cell culture process was accomplished with fed-batch bioreactor, and the downstream purification approach incorporated the combination of filtrations and liquid chromatography to obtain ultra-pure EV product. As previously reported, utilizing our modEXO™ platform, we have developed engineered EVs for the delivery of drug payloads for the treatment of diverse diseases, such as cancer,^38^ Inflammatory bowel disease,^39^ etc.

## 3. Material and Method Materials

Capto Core 700 was purchased from Cytiva. The SDA-100 chromatography system was purchased from Sepure Instruments. All buffers used in this study were prepared using ultrapure water (18.2 MΩ cm) from a Millipore Milli-Q water purification system.

Sodium bicarbonate, Formic acid, and TEAB were purchased from Sigma-Aldrich. Urea, Dithiothreitol (DTT), and iodoacetamide (IAM) were obtained from Amresco. Bovine serum albumin was sourced from Thermo Scientific. Trypsin was obtained from Promega. Ziptip was purchased from Millipore. Acetonitrile was sourced from J.T.Baker. Ammonia solution was obtained from Wako Pure Chemical Industries Ltd. Vials and caps were obtained from Thermo.

## Methods

### Cell culture

Cell density (VCD) and cell viability were assessed using a cell counter (Countleader FL 1000, Applitech); pH, pCO_2_, and pO_2_ were analyzed using a blood gas analyzer (RAPIDLab 348EX, SIEMENS); glucose, lactate, and ammonium ions were quantified using Cedex Bio (Roche); and osmotic pressure was determined using a freezing point osmometer (Gonotec). After cell culture, the supernatant was centrifuged at 4500-6000rpm for 30 minutes to remove cells. Subsequently, the supernatant was filtered through a 0.22μm sterile filter membrane for following up liquid chromatography procedure. Nanoparticle tracking analysis technology (ZetaView) was employed to measure the concentration and particle size of extracellular vesicles.

### Bind-elute size exclusion chromatography

Capto Core 700 chromatography column was equilibrated and the sample was loaded at 30∼50% of the column volume (CV), and the flow-through containing the sample was collected.

### Anion Exchange Chromatography

Sartobind® Q Nano membrane was equilibrated and loaded with a density of E12 P/ml. After washing with equilibration buffer, elution was performed.

### TEM (Transmission Electron Microscopy)

EVs were carefully deposited onto a copper grid (200 mesh) pre-coated with a carbon film and glow-discharged. Negative staining was performed by incubating the EVs with a 2% uranyl acetate solution at room temperature for 1 minute. After staining, excess stain was swiftly washed away with distilled water. The grids were then allowed to air-dry before being subjected to imaging using a Tecnai G2 transmission electron microscope (Thermo FEI, 120 kV).

### NTA (Nanoparticle Tracking Analysis)

EVs suspended in PBS solution were diluted in a range from 1E7 to 5E8/ml and introduced directly into the ZetaView Nano Particle Tracking Analyzer (ParticleMetrix, PMX120-Z).

### WB (Western Blot)

The western blotting procedure began with treating the sample with SDS and DTT solution at 95°C for 10 minutes to denature proteins. Gel electrophoresis was then conducted using a pre-cast SDS polyacrylamide gel in Tris-MOPs-SDS running buffer (GenScript) on a Tanon EPS-600 electrophoresis system. Subsequently, the gel was carefully removed and immersed in protein transfer buffer before being transferred onto a nitrocellulose membrane. The membrane was then subjected to staining with primary antibodies, including CD9 (PTG, 60232-1-Ig), CD63 (PTG, 25682-1-AP), CD81 (PTG, 66866-1-Ig), TSG101 (BD, 612696), beta Actin (ThermoFisher, MA5-16410), followed by secondary antibodies, including HRP Goat Anti-Rabbit IgG (H+L) (ABclonal, AS014), and HRP Goat Anti-Mouse IgG (H+L) (ABclonal, AS003). The luminescence readout was conducted using a Tanon-5200 Chemiluminescent Imaging System.

### Size Exclusion High-Performance Liquid Chromatography (SE-HPLC)

The SE-HPLC analysis was performed on an Agilent 1260 Infinity II HPLC system. The flow rate was set at 0.5 ml/min. Isocratic elution was employed throughout the analysis.

### Liquid Chromatography - Mass Spectrometry (LC-MS) proteomics analysis

The exosome was lysed and the protein was extracted. The protein sample was treated with 5mM DTT and incubated at 37°C for 1 hour, then returned to room temperature. Iodoacetamide was added to the mixture to a final concentration of 10 mM and incubate in the dark at room temperature for 45 minutes. The sample was diluted 4 times with 25 mM ammonium bicarbonate. Trypsin was added to the sample at a protein to enzyme ratio of 50:1 and incubate overnight at 37°C. The next day, the pH of the mixture was adjusted to less than 3 with formic acid to terminate the enzymatic digestion. The sample was desalted using a C18 desalting column, and then lyophilized.

For LC-MS analysis, mobile phase A (100% water, 0.1% formic acid) and mobile phase B (80% acetonitrile, 0.1% formic acid) were prepared. The lyophilized sample powder was dissolved in 10μL of mobile phase A, centrifuged at 14000×g for 20 min at 4°C, and 1 μg of the supernatant was used for liquid chromatography-mass spectrometry (LC-MS) analysis. A Q-Exactive HF-X mass spectrometer with a Nanospray Flex™ (NSI) ion source was used with the 2.4kV ion spray voltage, and 275°C ion transfer tube temperature. Mass spectrometry is performed in data-dependent acquisition mode, with a full scan range of m/z 350-1500, primary mass resolution set to 120,000 (at 200 m/z), and AGC set to 3×106. The maximum injection time for the C-trap is 80 ms. The top 40 ions from the full scan are selected for high-energy collision-induced dissociation (HCD) fragmentation in the second stage of mass spectrometry. Secondary mass resolution is set to 15,000 (at 200 m/z), with AGC set to 5×104 and a maximum injection time of 45 ms. The collision energy for peptide fragmentation is set to 27%.

## 4. Results and Discussion

Extracellular vesicles (EVs) have gained considerable attention in recent years due to their potential applications in various fields, including therapeutics, diagnostics, and drug delivery. However, the large-scale manufacture of EVs presents remains challenging. As shown in Figure 1, we introduce our bulk EV manufacture process in the aspect of upstream cell culture process development, downstream purification process development, and analytical development.

**Figure 1.**
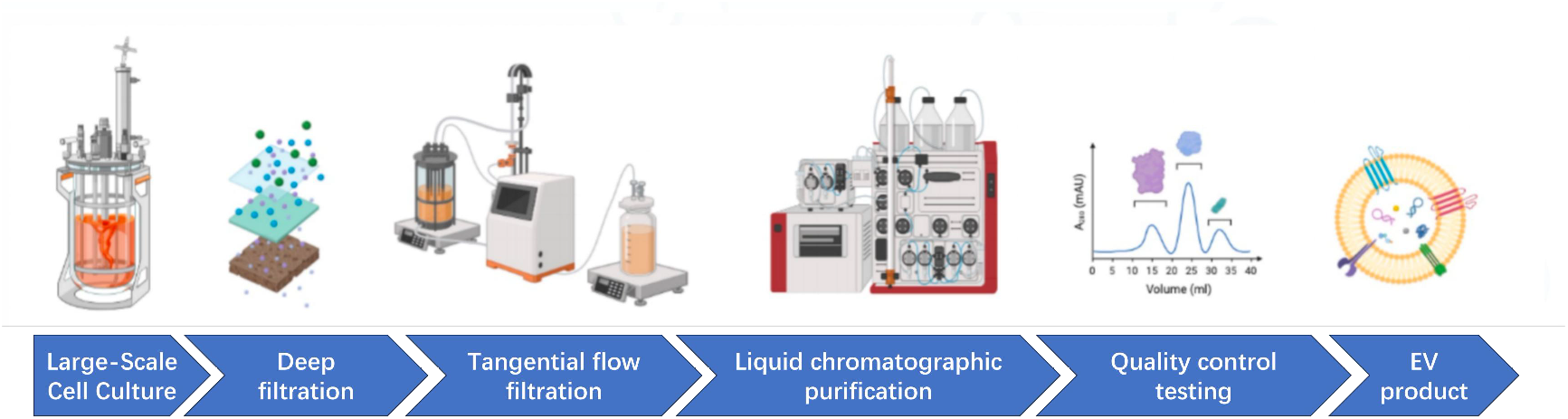
Schematic illustration of the principles of the large scale EV manufacture process workflow.

### Upstream process

To study the optimal condition for upstream cell culture process, three strains of suspended cells were utilized for the preparation of extracellular vesicles: cell line 1 (Expi293F engineered modification type), cell line 2 (Expi293F wild type), and cell line 3 (HEK293SF wild type). All three cell strains were resuscitated and passaged in serum-free culture medium, with culturing carried out at 37°C, 5% CO_2_, and expansion to N-1 generation at 100 rpm.

In the N phase, a serum-free culture medium was utilized for fed-batch cell culture. The cell culture period lasted 13 days with the addition of Feed Medium from Day 3. The cell culture temperature was maintained at 36.5°C, dissolved oxygen was controlled at 40%, and pH was regulated between 6.7 and 7.3 by the manipulation of CO_2_ gas and Na_2_CO_3_.

To monitor the cell growth metabolism as well as the quality detection, daily sampling of the supernatant from the bioreactor culture was carried out to measure cell density, viability, and biochemical parameters. Figure 2(A-F) illustrates the cell growth metabolism and EV expression during the fed-batch cell culture process at reactor scale for three cell strains. Through fed-batch cell culture, the three cell strains achieved densities of 38-50E6/mL after a 13-day cell culture period, with the modified Expi293F cell strain exhibiting the highest peak cell density (PK-VCD) reaching 48E6/mL (Figure 2A). At harvest on Day 13, the viability of the three cell strains remained above 90%, indicating overall favorable cell conditions (Figure 2B). Analysis of cell metabolism revealed similar trends among the three strains. Lactate accumulation occurred during the early growth phase until reaching the peak value, followed by a consumption phase, with peak values below 2g/L at harvest (Day 13), and lower lactate accumulation compared to HEK293 cells (Figure 2C). Ammonium ion concentration showed continuous accumulation during the early growth phase (D0-D4), followed by a brief consumption trend in the mid-growth phase before accumulating again in the late growth phase. HEK293 exhibited the highest overall ammonium ion accumulation, nearing 5mM at harvest, while the other two strains remained below 4mM (Figure 2D). Throughout the cell culture period, osmotic pressure showed an overall increasing trend due to the addition of feed culture medium, increased nutrient levels, and metabolic waste production. At harvest, the osmotic pressure of the three cell strains was close to each other, below 370 mOsm/kg (Figure 2E). Figure 2F shows the expression of EVs harvested from Day 10 to Day 13. Cell-line 2 and cell-line 3, two wild-type cell strains, exhibited similar trends in EV expression, with no significant increase from Day 10 to Day 13, and a slight decrease in expression at harvest. However, Cell-line 3, engineered with specific genetic modifications, showed increasing EV expression with prolonged cell culture time, significantly higher than the wild-type strains. At harvest, the EV expression level even reached twice that of the other two strains.

**Figure 2.**
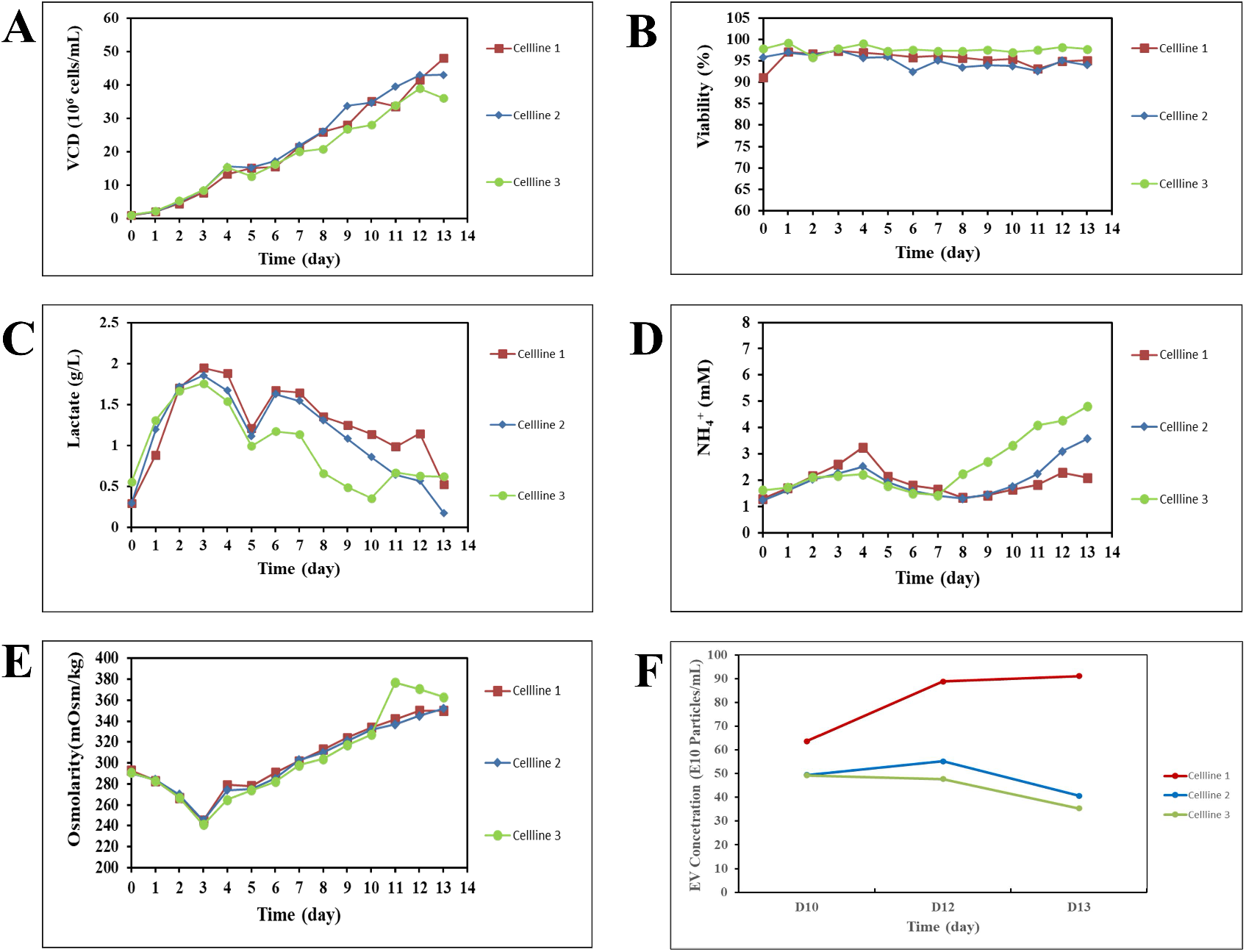
Cell culture monitoring for the upstream process development. (A) Cell density. (B) Cell viability. (C) Lactate. (D) Ammonium ion. (E) Osmolarity. (F) EV concentration.

Considering the comprehensive results of cell growth metabolism and EV expression, the use of fed-batch cell culture for the production of extracellular vesicles from HEK293 cell strains can achieve high levels of cell density and maintain high cell viability, thereby achieving high-yield extracellular vesicles. Additionally, fed-batch cell culture offers flexibility in production scale, allowing for the selection of different amplification scales based on sample volume requirements.

### Downstream Process

Extracellular vesicles (EVs) are characterized by their large size (50∼200nm) and abundant negative surface charge. Conventional chromatographic methods for bioproducts include affinity chromatography (AC), ion exchange chromatography, hydrophobic interaction chromatography, and size exclusion chromatography (SEC). Although EVs bear characteristic surface proteins, there are currently no affinity chromatography matrices available on the market that meet the requirements of chromatography (reproducibility, durability), thus rendering affinity chromatography unsuitable for EV capture. Based on the characteristics of EVs, we can separate smaller non-vesicular proteins and nucleic acids through size exclusion chromatography, followed by the removal of larger particles and other impurities through anion and cation exchange chromatography.

#### Bind-elute size exclusion chromatography

Capto Core 700 resin consists of a core matrix with activated ligands and an outer inert shell. During chromatographic flow-through, larger molecules are collected while smaller impurities bind to the internal ligands. The BE-SEC chromatogram shows good resolution between EVs and impurities, achieving baseline separation, as shown in Figure 3A. BE-SEC chromatography removes over 80% of protein impurities, with EV recovery exceeding 80%. TEM and SEC-HPLC testing results (Figure 4A, Figure 5A) support these findings, indicating the presence of larger protein impurities in the sample.

**Figure 3.**
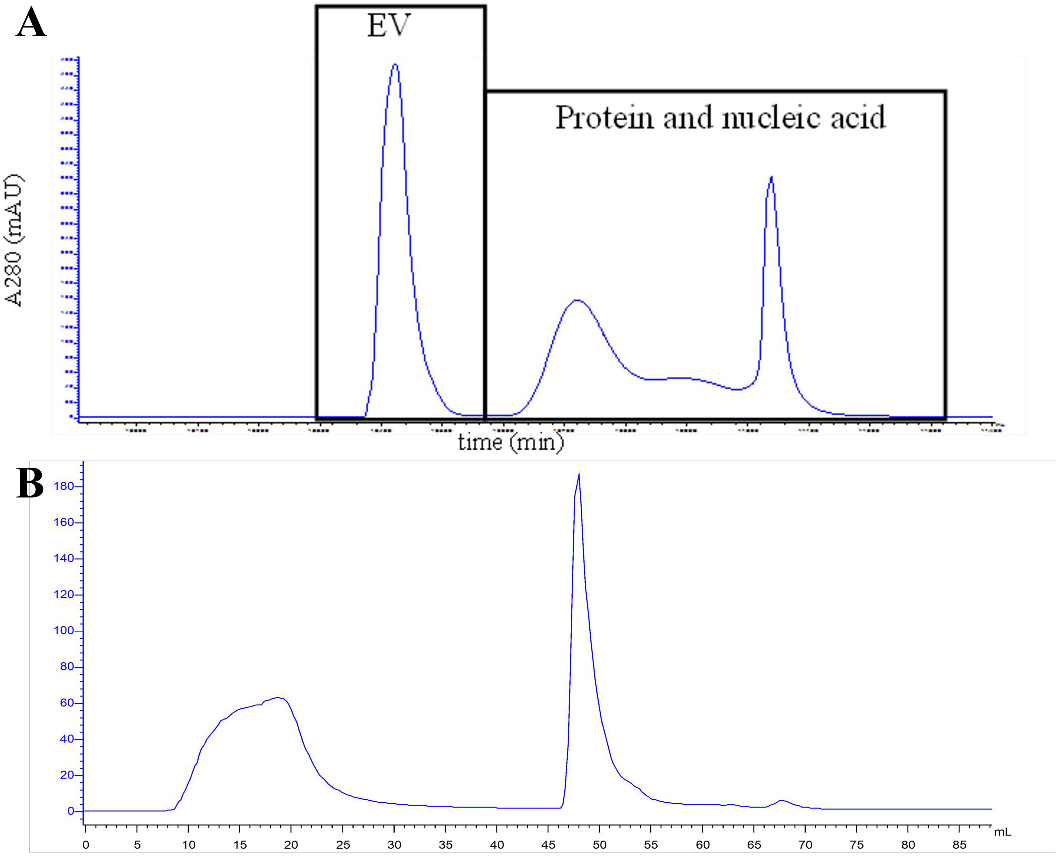
Chromatographic EV purification. (1) BE-SEC chromatogram. The first section is collected for detection, and the second section is CIP cleaning impurities, mainly proteins and nucleic acids. (B) AEX chromatogram.

**Figure 4.**
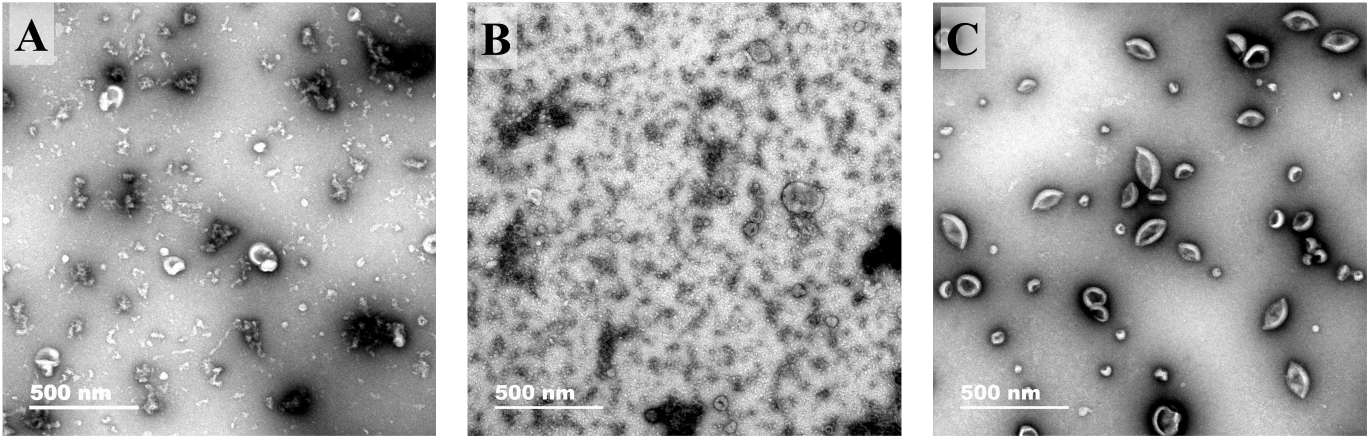
Electron Microscopy Image for EV samples. (A) EV sample collected from BE-SEC chromatography. (B) EV sample from AEX flow-through. (C) Sample from AEX elution.

**Figure 5.**
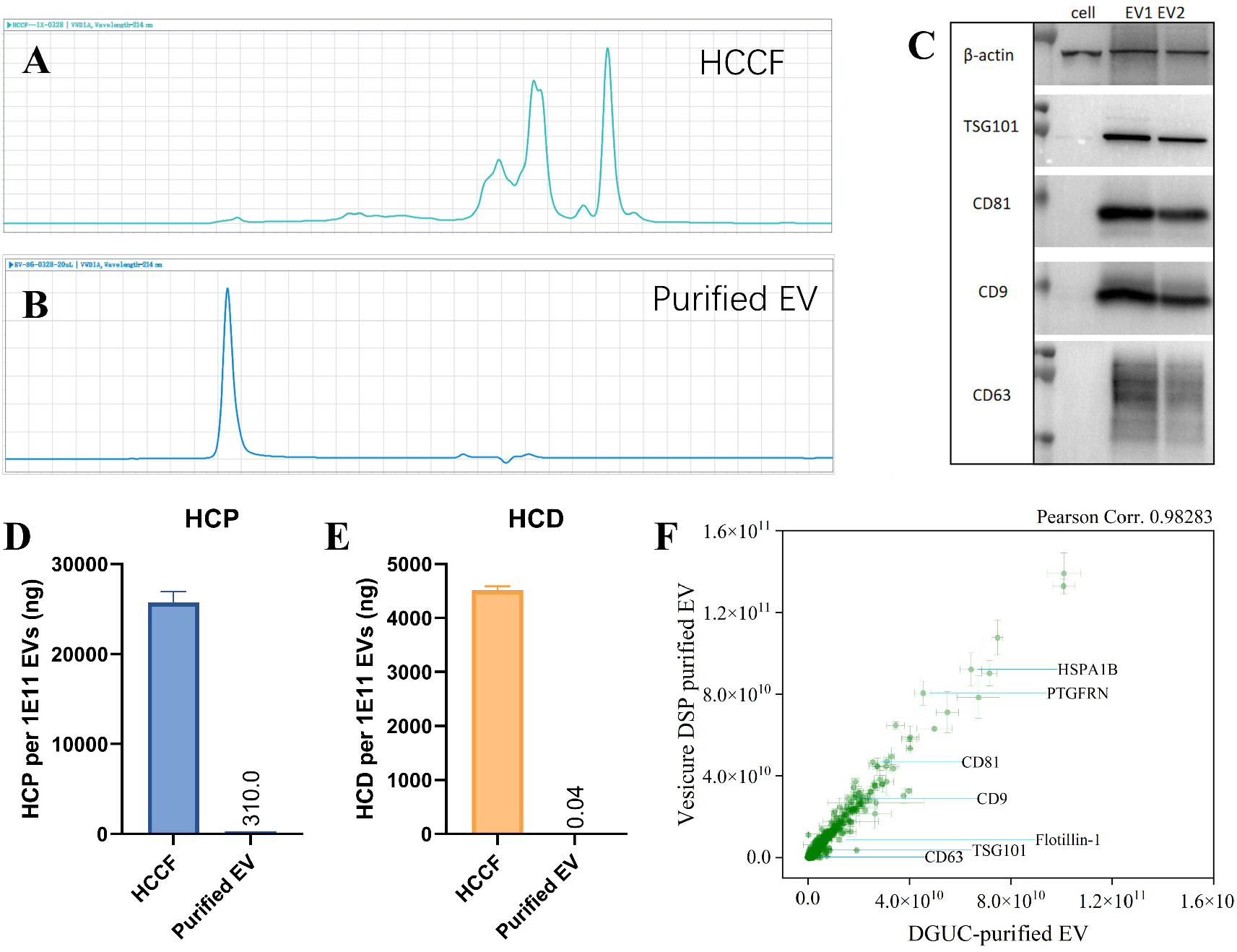
Analytical characterizations of purified EVs. (A) SEC-HPLC chromatograph of harvested cell culture fluid (HCCF). (B) SEC-HPLC chromatograph of EV product that are purified with VesiCURE downstream liquid chromatographic strategy. (C) An image of Western blot (WB) analysis targeting on EV markers. EV1 is the HEK293 EVs that are purified with density gradient ultracentrifugation. EV2 is the HEK293 EVs that are purified with VesiCURE downstream liquid chromatographic strategy. (D) Quantification of HEK293 host cell protein (HCP) on purified EV product. (E) Quantification of HEK293 host cell residual DNA (HCD) on purified EV product. (F) Proteomics data comparison between gradient-density ultracentrifugation purified EV and VesiCURE downstream chromatographic purified EV.

#### Anion Exchange Chromatography

During the culture process of HEK293 cells, a large amount of proteins is produced, varying in size, quantity, and properties. However, based on their different surface net charges, they can be generally classified into two categories: acidic proteins and basic proteins. Theoretically, we can remove acidic proteins through anion exchange chromatography. As seen from Figure 3B, a considerable amount of acidic proteins flow through, although the presence of EVs can also be observed. TEM result (Figure 4C) suggests that there is a noticeable impurity clearance, but still, some impurities remain. We have reasons to believe that these impurity proteins are similar in nature to EVs, and thus cannot be removed through a single purification step.

#### Process Scale-Up and Confirmation

We validated the stability of the process through purification at a scale of 3 batches of 15L each, ensuring consistent product quality. The average total recovery rate of the 3 batches was consistent with the small-scale results (Table 1), indicating relative stability and high reproducibility of the process. Next, we evaluated the differences in EVs obtained through chromatography and density gradient purification, focusing primarily on surface marker proteins, SEC purity, TEM, and mass spectrometry.

**Table 1.**
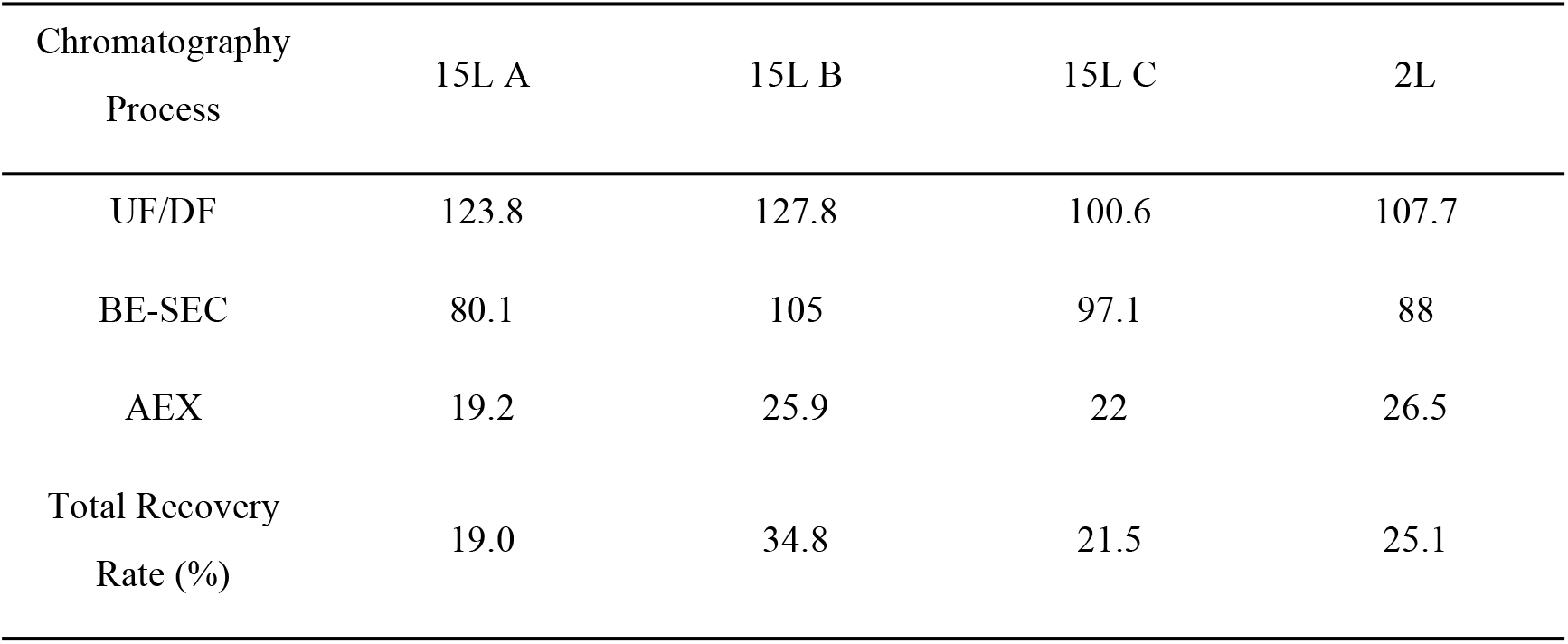
Recovery Rate of Chromatographic Purification.

### Analytical characterization

Analytical measurements play a pivotal role in ensuring the reliability, safety, and efficacy of extracellular vesicle production. We developed a suite of analytical methods across different aspects of EV characterization to comprehensively assess their quality, including (1) general characteristics, (2)identity verification, (3)contents analysis, (4)purity evaluations, (5)impurity assessments.

General characteristics includes visual inspection as the first step, examining the appearance, color, and clarity of the EV suspension. Instruments such as a pH meter and osmometer can be utilized to determine the pH and osmolality of the solution, respectively. Additionally, nanoparticle tracking analysis (NTA) were utilized to measure the diameter of EV particles, providing information about their size distribution as well as the Zeta potential information.

Identity verification involves detailed morphological examination using transmission electron microscopy (TEM) to visualize the characteristic cup-shaped morphology of EVs. As shown in Figure 4, the morphology of EVs and even the impurities can clearly be seen in TEM images. Immunoblotting (Western blotting) is a widely used technique to detect specific EV markers such as CD63, CD9, and CD81, providing evidence of their identity. As shown in Figure 5C, the EV marker proteins are detected highly abundant in our purified EV samples.

Contents analysis encompasses various techniques to quantify the content of EV preparations. We measured particle number using size exclusion chromatography (SEC) HPLC method and nanoparticle tracking analysis (data not shown). Total protein content was quantified using colorimetric assays such as the bicinchoninic acid (BCA) assay. Additionally, we performed LC-MS proteomic analysis on our EV product, profiling the protein expression of EVs from virous cell culture and purification approaches. Figure 5F shows a comparison of protein expression distribution between EVs that were purified with Density-gradient Ultracentrifugation and VesiCURE downstream chromatographic purification process, suggesting that minimal difference was present between these two samples, reflecting the ultra-high purity of our EV product obtained with our large-scale purification strategy.

Purity evaluations primarily rely on SEC-HPLC method, which highlights contaminants and macromolecular impurities based on size exclusion principles. EVs flow through the SEC column with a short retention time, while the impurities, if any, may elute out with a different retention time. As shown in Figure 5A, numerous impurities were observed in the Harvested Cell Culture Fluid (HCCF) sample, while the EV peak was barely seen. After purification, only the EV peak presented in the chromatogram, without any impurity peak but the buffer peak.

Impurity assessments involve detecting host cell proteins and DNA, which may contaminate EV preparations. We employed commercial Enzyme-linked immunosorbent assays (ELISA) kit (Yeasen, China) for host cell protein (HCP) quantification, and commercial quantitative polymerase chain reaction (qPCR) kit (Yeasen, China) for host cell residual DNA (HCD) detection. As shown in Figure 5D and 5E, over 99% clearance of HCP and 99.99% clearance of HCD were achieved with our downstream chromatographic purification strategy, ensuring that our EV product is free from host cell-derived impurities, minimizing potential immunogenicity risks.

## 5. Conclusion

In summary, our research has provided a comprehensive exploration of the upstream cell culture process for the production of extracellular vesicles (EVs) and the downstream purification strategies to obtain high-quality EV products. Through meticulous optimization of cell culture conditions and utilization of fed-batch cell culture techniques, we have demonstrated the ability to achieve high cell densities and maintain exceptional cell viability, resulting in high-yield EV production. Moreover, our downstream purification methods, including bind-elute size exclusion chromatography (BE-SEC) and anion exchange chromatography (AEX), have proven effective in separating and purifying EVs from cellular debris and other impurities.

Analytical characterization further confirms the purity and identity of the EVs obtained, ensuring their suitability for various applications in therapeutics, diagnostics, and drug delivery. Overall, our study provides valuable insights and methodologies for the scalable production and purification of EVs, paving the way for their widespread utilization in biomedical research and clinical applications.

## References

(1) Whitford, W.; Guterstam, P. Exosome manufacturing status. Futuremedicinalchemistry 2019, 11(10), 1225–1236.

(2) Simons, M.; Raposo, G. Exosomes–vesicular carriers for intercellular communication. Currentopinionincellbiology 2009, 21(4), 575–581.

(3) Malm, T.; Loppi, S.; Kanninen, K. M. Exosomes in Alzheimer’s disease. Neurochemistry international 2016, 97, 193–199.

(4) Van Giau, V.; An, S. S. A. Emergence of exosomal miRNAs as a diagnostic biomarker for Alzheimer’s disease. Journaloftheneurologicalsciences 2016, 360, 141–152.

(5) Xu, Z.; Zeng, S.; Gong, Z.; Yan, Y. Exosome-based immunotherapy: a promising approach for cancer treatment. Molecular cancer 2020, 19, 1–16.

(6) Biosciences, M. b. C. Exosome cancer diagnostic reaches market. Nat.Biotechnol 2016, 34, 359.

(7) Wang, J.; Zheng, Y.; Zhao, M. Exosome-based cancer therapy: implication for targeting cancer stem cells. Frontiersinpharmacology 2017, 7, 533.

(8) Zhao, W.; Zheng, X.-L.; Zhao, S.-P. Exosome and its roles in cardiovascular diseases. HeartFailureReviews 2015, 20, 337–348.

(9) Kalluri, R.; LeBleu, V. S. The biology, function, and biomedical applications of exosomes. Science 2020, 367(6478), eaau6977.

(10) Wu, M.; Ouyang, Y.; Wang, Z.; Zhang, R.; Huang, P.-H.; Chen, C.; Li, H.; Li, P.; Quinn, D.; Dao, M. Isolation of exosomes from whole blood by integrating acoustics and microfluidics. Proceedings of the National Academy of Sciences 2017, 114 (40), 10584–10589.

(11) Park, O.; Choi, E. S.; Yu, G.; Kim, J. Y.; Kang, Y. Y.; Jung, H.; Mok, H. Efficient delivery of tyrosinase related protein-2 (TRP2) peptides to lymph nodes using serum-derived exosomes. Macromolecular Bioscience 2018, 18(12), 1800301.

(12) Baek, G.; Choi, H.; Kim, Y.; Lee, H.-C.; Choi, C. Mesenchymal stem cell-derived extracellular vesicles as therapeutics and as a drug delivery platform. Stemcellstranslational medicine 2019, 8(9), 880–886.

(13) Upadhya, R.; Madhu, L. N.; Attaluri, S.; Gitaí, D. L. G.; Pinson, M. R.; Kodali, M.; Shetty, G.; Zanirati, G.; Kumar, S.; Shuai, B. Extracellular vesicles from human iPSC-derived neural stem cells: miRNA and protein signatures, and anti-inflammatory and neurogenic properties. Journalofextracellularvesicles 2020, 9(1), 1809064.

(14) Lai, R. C.; Yeo, R. W. Y.; Padmanabhan, J.; Choo, A.; De Kleijn, D. P.; Lim, S. K. Isolation and characterization of exosome from human embryonic stem cell-derived C-Myc-immortalized mesenchymal stem cells. Mesenchymal Stem Cells: Methods and Protocols 2016, 477–494.

(15) Estes, S.; Konstantinov, K.; Young, J. D. Manufactured extracellular vesicles as human therapeutics: challenges, advances, and opportunities. CurrentOpinioninBiotechnology 2022, 77, 102776.

(16) Deatherage, B. L.; Cookson, B. T. Membrane vesicle release in bacteria, eukaryotes, and archaea: a conserved yet underappreciated aspect of microbial life. Infectionandimmunity 2012, 80(6), 1948–1957.

(17) Adriano, B.; Cotto, N. M.; Chauhan, N.; Jaggi, M.; Chauhan, S. C.; Yallapu, M. M. Milk exosomes: Nature’s abundant nanoplatform for theranostic applications. Bioactive materials 2021, 6(8), 2479–2490.

(18) Marsh, S. R.; Pridham, K. J.; Jourdan, J.; Gourdie, R. G. Novel protocols for scalable production of high quality purified small extracellular vesicles from bovine milk. Nanotheranostics 2021, 5(4), 488.

(19) Kim, J.; Li, S.; Zhang, S.; Wang, J. Plant-derived exosome-like nanoparticles and their therapeutic activities. AsianJournalofPharmaceuticalSciences 2022, 17(1), 53–69.

(20) Picciotto, S.; Barone, M. E.; Fierli, D.; Aranyos, A.; Adamo, G.; Božič, D.; Romancino, D. P.; Stanly, C.; Parkes, R.; Morsbach, S. Isolation of extracellular vesicles from microalgae: Towards the production of sustainable and natural nanocarriers of bioactive compounds. Biomaterials Science 2021, 9(8), 2917–2930.

(21) Abaandou, L.; Quan, D.; Shiloach, J. Affecting HEK293 Cell Growth and Production Performance by Modifying the Expression of Specific Genes. Cells 2021, 10 (7). DOI: 10.3390/cells10071667 From NLM.

(22) Colao, I. L.; Corteling, R.; Bracewell, D.; Wall, I. Manufacturing exosomes: a promising therapeutic platform. Trendsinmolecularmedicine 2018, 24(3), 242–256.

(23) Ahn, S.-H.; Ryu, S.-W.; Choi, H.; You, S.; Park, J.; Choi, C. Manufacturing therapeutic exosomes: from bench to industry. MoleculesandCells 2022, 45(5), 284.

(24) Dhondt, B.; Geeurickx, E.; Tulkens, J.; Van Deun, J.; Vergauwen, G.; Lippens, L.; Miinalainen, I.; Rappu, P.; Heino, J.; Ost, P. Unravelling the proteomic landscape of extracellular vesicles in prostate cancer by density-based fractionation of urine. Journalof extracellularvesicles 2020, 9(1), 1736935.

(25) Yang, D.; Zhang, W.; Zhang, H.; Zhang, F.; Chen, L.; Ma, L.; Larcher, L. M.; Chen, S.; Liu, N.; Zhao, Q. Progress, opportunity, and perspective on exosome isolation-efforts for efficient exosome-based theranostics. Theranostics 2020, 10(8), 3684.

(26) Muller, L.; Hong, C.-S.; Stolz, D. B.; Watkins, S. C.; Whiteside, T. L. Isolation of biologically-active exosomes from human plasma. Journalofimmunologicalmethods 2014, 411, 55–65.

(27) Livshits, M. A.; Khomyakova, E.; Evtushenko, E. G.; Lazarev, V. N.; Kulemin, N. A.; Semina, S. E.; Generozov, E. V.; Govorun, V. M. Isolation of exosomes by differential centrifugation: Theoretical analysis of a commonly used protocol. Scientificreports 2015, 5 (1), 17319.

(28) Paolini, L.; Zendrini, A.; Noto, G. D.; Busatto, S.; Lottini, E.; Radeghieri, A.; Dossi, A.; Caneschi, A.; Ricotta, D.; Bergese, P. Residual matrix from different separation techniques impacts exosome biological activity. Scientificreports 2016, 6(1), 23550.

(29) Tatischeff, I.; Larquet, E.; Falcón-Pérez, J. M.; Turpin, P.-Y.; Kruglik, S. G. Fast characterisation of cell-derived extracellular vesicles by nanoparticles tracking analysis, cryo-electron microscopy, and Raman tweezers microspectroscopy. Journalofextracellular vesicles 2012, 1(1), 19179.

(30) Li, P.; Kaslan, M.; Lee, S. H.; Yao, J.; Gao, Z. Progress in exosome isolation techniques. Theranostics 2017, 7(3), 789.

(31) Soares Martins, T.; Catita, J.; Martins Rosa, I.; AB da Cruz e Silva, O.; Henriques, A. G. Exosome isolation from distinct biofluids using precipitation and column-based approaches. PloSone 2018, 13(6), e0198820.

(32) Lim, C. Z.; Zhang, L.; Zhang, Y.; Sundah, N. R.; Shao, H. New sensors for extracellular vesicles: insights on constituent and associated biomarkers. ACSsensors 2019, 5(1), 4–12.

(33) Gámez-Valero, A.; Monguió-Tortajada, M.; Carreras-Planella, L.; Franquesa, M. l.; Beyer, K.; Borràs, F. E. Size-Exclusion Chromatography-based isolation minimally alters Extracellular Vesicles’ characteristics compared to precipitating agents. Scientific reports 2016, 6(1), 33641.

(34) Heath, N.; Grant, L.; De Oliveira, T. M.; Rowlinson, R.; Osteikoetxea, X.; Dekker, N.; Overman, R. Rapid isolation and enrichment of extracellular vesicle preparations using anion exchange chromatography. Scientificreports 2018, 8(1), 5730.

(35) Noyes, A.; Doherty, M.; Ellis, K.; Bourdeau, R.; Desanty, K. Process for preparing extracellular vesicles. Google Patents: 2022.

(36) de Fougerolles, T.; Evox’s, C. Engineering exosomes to create transformational drugs.

(37) Wiklander, O.; Gorgens, A. Affinity purification of engineered extracellular vesicles. Google Patents: 2021.

(38) Zhai, Y.; Niu, S.; Qiu, T.; Yi, Y.; Yan, Y.; Hu, R.; He, X.; Xu, K. Therapeutic targeting of KRAS mutation-driven tumorigenesis by extracellular vesicles loaded with small interfering RNA. bioRxiv 2024, 2024.2001. 2017.576015.

(39) Zhai, Y.; Liu, G.; Cui, C.; Yi, Y.; Yan, Y.; Zhang, L.; Wang, W.; He, X.; Xu, K. Mesenchymal stem cell-derived extracellular vesicles attenuate symptoms in dextran sodium sulfate-induced ulcerative colitis mouse model. bioRxiv 2024, 2024.2002. 2001.578325.

